# FRMD8 promotes inflammatory and growth factor signalling by stabilising the iRhom/ADAM17 sheddase complex

**DOI:** 10.1101/255802

**Authors:** Ulrike Künzel, Adam G. Grieve, Yao Meng, Sally A. Cowley, Matthew Freeman

**Affiliations:** Sir William Dunn School of Pathology, University of Oxford, South Parks Road, Oxford OX1 3RE

## Abstract

Many intercellular signals are synthesised as transmembrane precursors that are released by proteolytic cleavage (‘shedding’) from the cell surface. ADAM17, a membrane-tethered metalloprotease, is the primary shedding enzyme responsible for the release of the inflammatory cytokine TNFα and several EGF receptor ligands. ADAM17 exists in complex with the rhomboid-like iRhom proteins, which act as cofactors that regulate ADAM17 substrate shedding. Here we report that the poorly characterised FERM domain-containing protein FRMD8 is a new component of iRhom2/ADAM17 sheddase complex. FRMD8 binds to the cytoplasmic N-terminus of iRhoms, and is necessary to stabilise the iRhoms and ADAM17 beyond the Golgi. In the absence of FRMD8, iRhom2 and ADAM17 are degraded via the endolysosomal pathway, resulting in the reduction of ADAM17-mediated shedding. We have confirmed the pathophysiological significance of FRMD8 in iPSC-derived human macrophages and mouse tissues, thus demonstrating its role in the regulated release of multiple cytokine and growth factor signals.

## Introduction

The cell surface protease ADAM17 (also called TACE) mediates the release of many important signalling molecules by ‘shedding’ their extracellular ligand domains from transmembrane precursors. A prominent example is ADAM17’s role in releasing the tumour necrosis factor alpha (TNFα) (Black et al., 1997, Moss et al., 1997), a primary cytokine involved in the inflammatory responses to infection and tissue damage (Kalliolias and Ivashkiv, 2016). In addition, ADAM17 is the principle sheddase of the epidermal growth factor (EGF) receptor ligands amphiregulin (AREG), transforming growth factor alpha (TGFα), heparin-binding EGF (HB-EGF), epigen, and epiregulin (Sahin et al., 2004, Sahin and Blobel, 2007). The control of ADAM17 activity has therefore been the focus of much fundamental and pharmaceutical research (reviewed in (Rose-John, 2013, Zunke and Rose-John, 2017)). We and others have previously reported that the rhomboid-like iRhom proteins have a specific and extensive regulatory relationship with ADAM17, to the extent that iRhoms can effectively be considered as regulatory subunits of the protease. iRhoms are members of a wider family of evolutionarily related multi-pass membrane proteins, called the rhomboid like superfamily (Freeman, 2014). The family is named after the rhomboids, intramembrane serine proteases that cleave substrate transmembrane domains (TMDs), but many members, including iRhoms, have lost protease activity during evolution.

iRhom1 and its paralogue iRhom2 (encoded by the genes *RHBDF1* and *RHBDF2*) show redundancy in regulating ADAM17 maturation, but differ in their tissue expression (Christova et al., 2013). Many cell types express both iRhoms, so the loss of one can be compensated by the other (Christova et al., 2013, Li et al., 2015). Macrophages are, however, an exception: iRhom1 is not expressed, so iRhom2 alone regulates ADAM17, and therefore TNFα inflammatory signalling in macrophages (Adrain et al., 2012, McIlwain et al., 2012, Issuree et al., 2013). iRhoms control ADAM17 activity in multiple ways. First, they bind to the catalytically immature pro-form of ADAM17 (proADAM17) in the endoplasmic reticulum (ER), and are required for its trafficking from the ER to the Golgi apparatus (Adrain et al., 2012, McIlwain et al., 2012). Once proADAM17 reaches the Golgi, it is matured by the removal of its inhibitory pro-domain by pro-protein convertases (Schlondorff et al., 2000, Endres et al., 2003) and is further trafficked to the plasma membrane. iRhoms have further regulatory functions beyond this step of ADAM17 maturation. Still bound to each other, iRhom2 prevents the lysosomal degradation of ADAM17 (Grieve et al., 2017). Later, iRhom2 controls the activation of ADAM17: phosphorylation of the iRhom2 cytoplasmic tail promotes the recruitment of 14-3-3 proteins, which promote shedding activity of ADAM17, thereby releasing TNFα from the cell surface in response to inflammatory triggers (Grieve et al., 2017, Cavadas et al., 2017). Finally, iRhoms are also reported to contribute to ADAM17 substrate specificity (Maretzky et al., 2013). This intimate regulatory role of iRhoms make them essential players in ADAM17-mediated signalling and thus new targets for manipulating inflammatory signalling. The significance of this potential is underlined by the fact that anti‐ TNFα therapies, used to treat rheumatoid arthritis and other inflammatory diseases, are currently the biggest grossing drugs in the world (Monaco et al., 2015).

Despite the role of the iRhom/ADAM17 shedding complex in controlling signalling, much is yet to be understood about the molecular mechanisms that control this inflammatory trigger. To identify the wider machinery by which iRhoms regulate ADAM17, we report here a proteomic screen to identify their binding partners. We have identified the poorly characterised FERM domain-containing protein 8 (FRMD8) as having a strong and specific interaction with the cytoplasmic region of iRhoms. The functional significance of this interaction is demonstrated by loss of FRMD8 causing a similar phenotype to iRhom deficiency in cells: loss of mature ADAM17 and severely reduced shedding of ADAM17 substrates from the cell surface. We show that loss of FRMD8 leads to lysosomal degradation of mature ADAM17 and iRhom2, indicating that although FRMD8 binds to iRhom2 throughout its entire life cycle, its main function is to stabilise iRhoms and ADAM17 once they reach the plasma membrane. Overall, our results imply that FRMD8 is an essential component of the inflammatory signalling machinery. To test this proposal *in vivo* we deleted the FRMD8 gene in human induced pluripotent stem cells (iPSCs) and differentiated them into macrophages. Consistent with our biochemical data, these mutant macrophages were defective in their ability to release TNFα in response to lipopolysaccharide (LPS) stimulation, demonstrating the pathophysiological importance of FRMD8 in the normal inflammatory response by human macrophages. The *in vivo* significance of FRMD8 in regulating the stability of the iRhom/ADAM17 shedding complex was further reinforced by our observation that mature ADAM17 and iRhom2 protein levels are strongly reduced in tissues of FRMD8-deficent mice.

## Results

### FRMD8 is a novel interaction partner of iRhom1 and iRhom2

To investigate the molecular mechanisms that underlie iRhom2 functions, we performed a mass spectrometry-based screen to identify new proteins that interact with human iRhom2. C-terminally tagged protein (iRhom2-3xHA) was stably expressed in human embryonic kidney (HEK) 293T cells and immunoprecipitated. The bead eluates containing immunoprecipitated iRhom2 and its interacting proteins were analysed by label-free mass spectrometry. As a negative control, we did the same analysis in parallel with 3xHA-tagged UNC93B1, an unrelated polytopic protein that, like iRhom2, is predominantly located in the ER (Koehn et al., 2007) (Fig. S1A). Quantitative protein abundance data from three biological replicates of Rhom2 and UNC93B1 co-immunoprecipitations were statistically analysed using the Perseus software platform (Tyanova et al., 2016). Validating the overall approach, we detected ADAM17, the known iRhom2 interacting protein (Adrain et al., 2012, McIlwain et al., 2012, Christova et al., 2013) as a statistically significant hit (Fig. 1A, S1B). Among other significant hits were several 14-3-3 proteins (eta, epsilon, gamma, sigma, theta, zeta/delta) and MAPK1/3 (Fig. S1B), which we have previously reported to participate in the regulation of inflammatory signalling by phosphorylation of iRhom2 (Grieve et al., 2017). The top hit by a long way, however, was FRMD8 (Fig. 1A, S1B), a poorly studied protein that has not previously been implicated in iRhom function, ADAM17 regulation, or growth factor or cytokine signalling.

**Figure 1.**
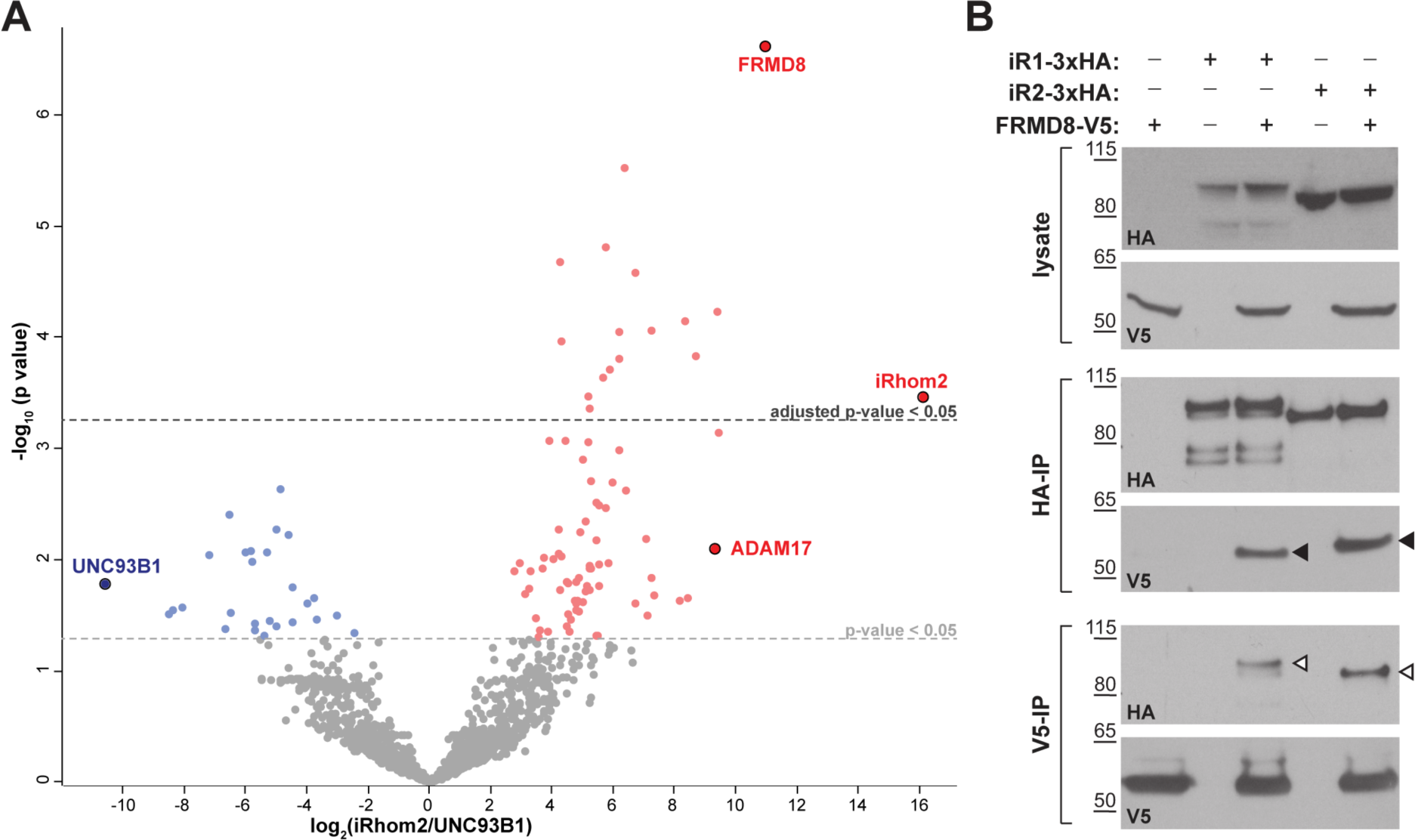
FRMD8 is a novel interaction partner of iRhom1 and iRhom2. **A** Volcano plot representing results from three iRhom2 co-immunoprecipitations. The fold change of label-free quantification values (in log2 ratio) was plotted against the p value (− log10 transformed). The grey dotted line indicates p-values < 0.05 (analysed with a two sample t-test). Benjamini-Hochberg correction was applied to adjust the p-value for multiple hypothesis testing (dark grey dotted line). **B** Lysates of HEK293T cells stably expressing human iRhom1-3xHA or iRhom2-3xHA transfected with human FRMD8-V5 (where indicated) were subjected to anti-HA and anti-V5 immunoprecipitation (HA-IP, V5-IP) and a western blot using anti-HA and anti-V5 antibodies was performed. Black arrowheads indicated the co-immunoprecipitated FRMD8-V5; white arrowheads indicated the co-immunoprecipitated iRhoms.

We confirmed the interaction between iRhom2 and FRMD8 by immunoprecipitation. C-terminally V5 tagged FRMD8 co-immunoprecipitated with either iRhom1-3xHA or iRhom2-3xHA (Fig. 1B). Conversely, we pulled down both iRhom1-3xHA and iRhom2-3xHA with an antibody against the V5 tag. Finally, we were also able to co-immunoprecipitate endogenous FRMD8 with iRhom2-3xHA (Fig. S1C). Together these results identify FRMD8 as a bona fide binding partner of iRhom1 and iRhom2 in human cells.

### FRMD8 is required for iRhom function

As its name indicates, FRMD8 is a FERM (4.1/ezrin/radixin/moesin) domain containing protein. It is predicted to be a soluble cytoplasmic protein, and the only report about its function describes it as binding to the Wnt accessory receptor low-density lipoprotein receptor-related protein 6 (LRP6), and negatively regulating Wnt signalling (Kategaya et al., 2009). To investigate the functional significance of FRMD8 binding to iRhoms, we examined the effects of loss of FRMD8 on iRhom function in HEK293T cells, using both siRNA and CRISPR/Cas9-mediated gene deletion (Fig. 2A, B). In both cases, loss of FRMD8 drastically reduced the protein levels of mature ADAM17 (Fig. 2A, B). This effect was specific to ADAM17, as the maturation of its closest homologue, ADAM10, was unaffected by loss of FRMD8 (Fig. 2B). Moreover, mature ADAM17 levels were rescued by expression of FRMD8-V5 in FRMD8 knockout HEK293T cells (Fig. 2C), confirming that the phenotype was caused by FRMD8 loss. Finally, in addition to this reduction of mature ADAM17 caused by FRMD8 loss, we found a striking loss of ADAM17, but not ADAM10, on the cell surface (Fig. 2D). These phenotypes partially phenocopy the loss of iRhoms (Christova et al., 2013, Grieve et al., 2017), consistent with FRMD8 being needed for iRhoms to act as positive regulators of ADAM17.

**Figure 2.**
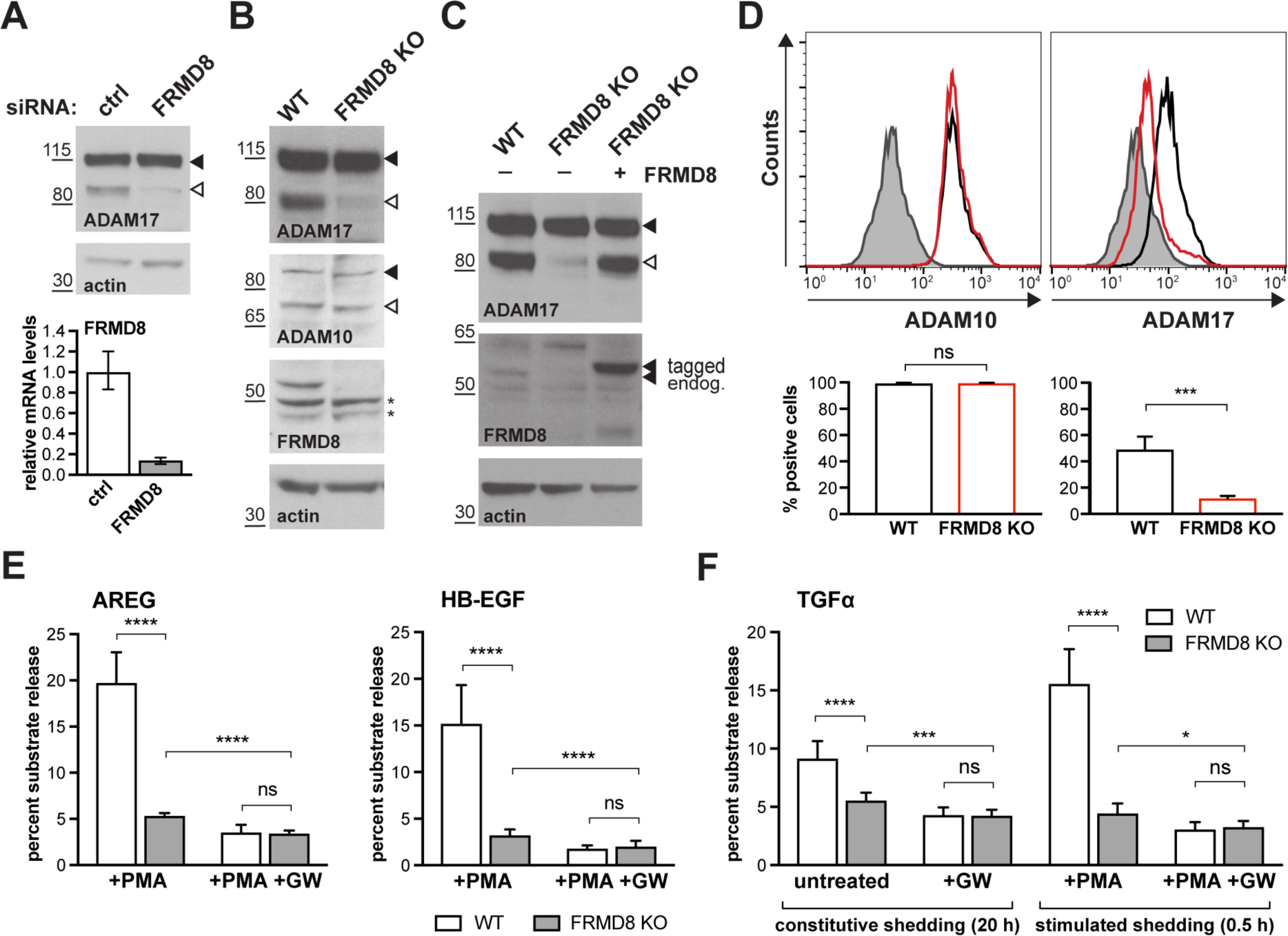
FRMD8 loss reduces mature ADAM17 levels and impairs ADAM17-dependent shedding activity. **A** ADAM17 levels were analysed in HEK293T cells transfected with non-targeting siRNA control pool (ctrl) or FRMD8 SMARTpool siRNA after western blotting with anti-ADAM17 and anti-actin staining. In this and subsequent figures, pro‐ and mature form of ADAM17 are indicated with black and white arrowheads, respectively. Lower panel: Knockdown efficiency of FRMD8 was analysed by TaqMan PCR. **B, C** Lysates from wild-type (WT) and FRMD8 knockout (KO) HEK293T cells were transiently transfected with FRMD8-V5 (where indicated) and immunoblotted for endogenous ADAM17, ADAM10, FRMD8 and actin using western blotting. Nonspecific bands are marked with an asterisk. **D** Cell surface levels of endogenous ADAM10 and ADAM17 were analysed in WT and FRMD8 KO HEK293T cells after stimulation with 200 nM PMA for 5 min. Unpermeabilised cells were stained on ice with ADAM10 and ADAM17 antibodies, or only with secondary antibody as a control (grey). The immunostaining was analysed by flow cytometry. The graph shown is one representative experiment out of four biological replicates. The percentage of positive cells was calculated for each experiment using FlowJo software. Statistical analysis was performed using an unpaired t-test. **E, F** WT and FRMD8 KO HEK293T cells were transiently transfected with alkaline phosphatase (AP)-tagged AREG, HB-EGF or TGFα, and then either incubated with 200 nM PMA, with 200 nM PMA and 1 μM GW (ADAM10/ADAM17 inhibitor), or with DMSO for 30 min. In addition, cells transfected with AP-TGFα were either left unstimulated for 20 h or incubated with GW for 20 h. AP activity was measured in supernatants and cell lysates. Each experiment was performed in biological triplicates. The results of three independent shedding experiments are shown. Statistical analysis was performed of using a Mann-Whitney test. ns = p-value > 0.05; * = p-value < 0.05; *** = p-value < 0.001; **** = p-value < 0.0001.

We also examined the consequences of loss of FRMD8 on ADAM17-dependent signalling. The shedding of alkaline phosphatase (AP)-tagged EGF receptor ligands AREG and HB-EGF, after stimulation with phorbol 12-myristate 13-acetate (PMA), were both substantially reduced in FRMD8 knockout cells (Fig. 2E). To exclude the possibility that the defect in FRMD8 knockout cells is an inability to respond to PMA, we measured both PMA-stimulated and unstimulated, constitutive shedding of AP-tagged TGFα, another major EGFR ligand. Again, FRMD8 knockout cells released significantly less AP-TGFα compared to wild-type cells, both after stimulation but also after 20 h of constitutive shedding (Fig. 2F), implying that mutant cells had fundamental defects in their ability to shed ADAM17 ligands, regardless of PMA stimulation. To demonstrate that the release of ligands was indeed caused by metalloprotease shedding and not simply an indication of leakage caused by cell death, we showed that it was sensitive to the ADAM10/17 inhibitor GW280264X (GW) (Fig.2E, F). Overall, as with ADAM17 maturation and localisation, the shedding defects in FRMD8-deficient cells resemble those caused by the loss of iRhoms.

### FRMD8 binds to the iRhom2 cytoplasmic N-terminus throughout the entire secretory pathway

As described above, iRhoms regulate ADAM17 function at multiple stages: from ER-to-Golgi trafficking, to the activation of the sheddase at the cell surface. To address where FRMD8 fits in this long-term relationship between iRhoms and ADAM17, we started by analysing where in the secretory pathway FRMD8 binds to iRhom2. Immunofluorescent staining of FRMD8-V5 with anti-V5 antibody confirmed what had been reported by Kategaya et al.(2009): the protein is detected in the cytoplasm and associated with the plasma membrane (Fig. 3A). When co-expressed with iRhom2-3xHA, we detected significant co-localisation between the proteins (Fig. 3A) further confirming the interaction of FRMD8 and iRhom2. Taking a more biochemical approach, immunoprecipitation of FRMD8 pulled down both immature and mature ADAM17 (Fig. 2B), indicating that there is a sufficiently stable tripartite complex between FRMD8, iRhom2 and ADAM17 to allow FRMD8 to co-immunoprecipitate ADAM17. Note that FRMD8 did not pull down ADAM17 in cells mutant for both iRhoms (Fig. S2A), implying that there is no direct link between them; instead both can bind simultaneously to iRhom2. Significantly, FRMD8 binds iRhom2 when in complex with immature ADAM17, which only exists in the ER and early Golgi apparatus, and also iRhom2 in complex with mature ADAM17, which exists in the *trans-*Golgi network and beyond; together these data imply that FRMD8 binds to iRhom2 throughout the secretory pathway.

**Figure 3.**
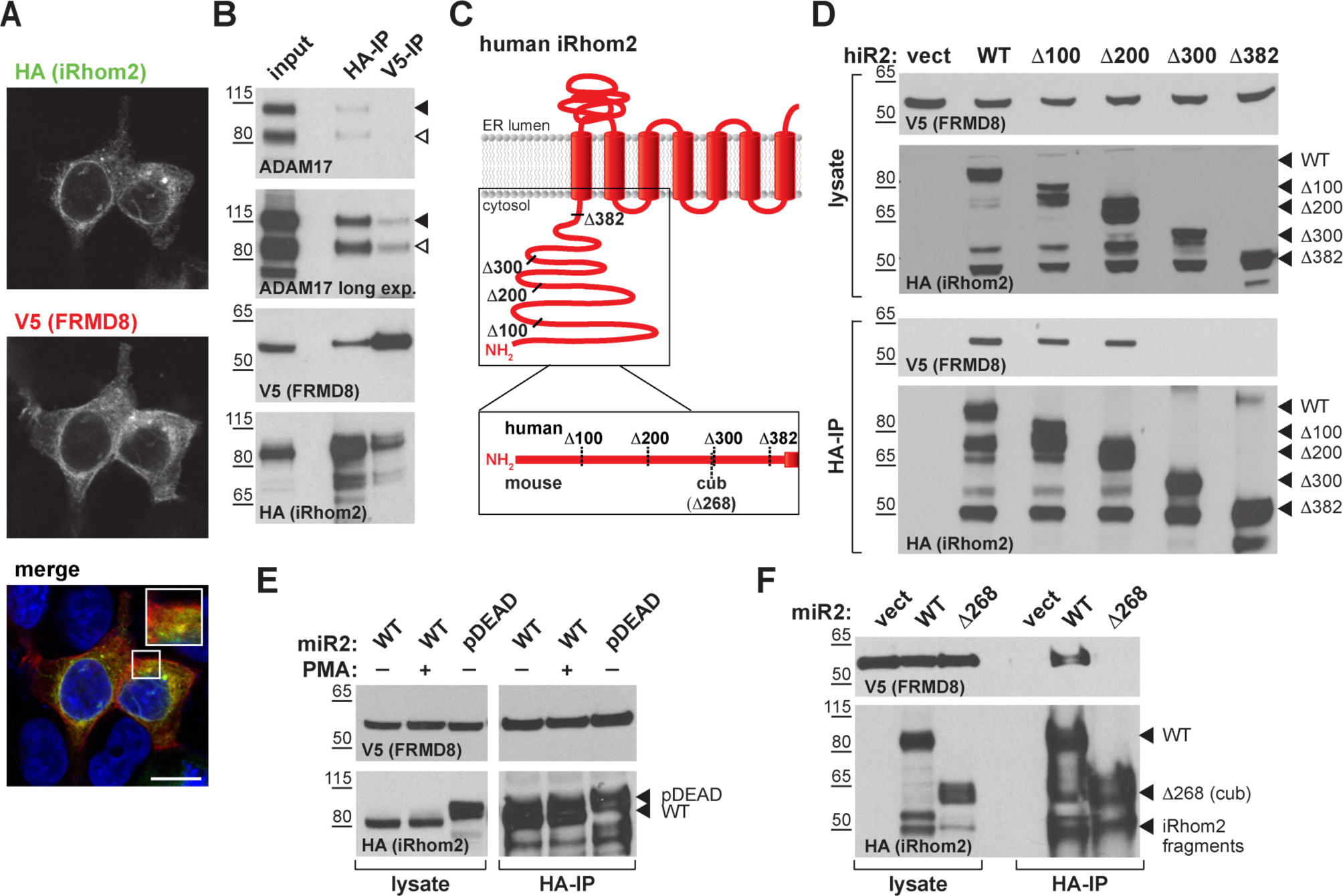
FRMD8 binds to the iRhom2 N-terminus throughout the entire secretory pathway. **A** MCF-7 cells were transfected with human iRhom2-3xHA and FRMD8-V5 and stained using 4′,6-diamidino-2-phenylindole (DAPI; blue; to mark nuclei), anti-HA and anti-V5 antibodies. Cells were imaged on a confocal fluorescent microscope. Scale bar: 10 μm. **B** Lysates from anti-HA and anti-V5 immunoprecipitation (HA-IP, V5-IP) of HEK293T cells co-expressing human iRhom2-3xHA and human FRMD8-V5 were immunoblotted for ADAM17, HA and V5. **C** Schematic representation of truncated human and mouse iRhom2 constructs used in (D) and (F). **D, E, F** Lysates and anti-HA immunoprecipitation (HA-IP) from HEK293T cells transiently transfected with FRMD8-V5 and either empty vector (vect), truncated human iRhom2-3xHA constructs (D), mouse iRhom2^WT^ (WT) and iRhom2^pDEAD^ (pDEAD) (E) or mouse iRhom2^WT^ and Rhom2^cub^ (Δ268) (F) were immunoblotted for V5 and HA. Where indicated cells have been stimulated with 200 nM PMA for 30 min.

As a cytoplasmic protein, FRMD8 was likely to bind to the only substantial cytoplasmic region of iRhom2, its N-terminus. We therefore made a set of iRhom2-3xHA N-terminal deletion constructs (Fig. 2C) to locate the binding site. Deletion of the first 200 amino acids in the N-terminus of iRhom2 did not disrupt FRMD8 binding, but no interaction was detected in mutants greater than ∆300 (Fig. 2D), implying that the region between 200 and 300 amino acids was necessary for FRMD8 binding. Although the N-terminal cytoplasmic tail of iRhom2 contains multiple regulatory phosphorylation sites (Fig. S2B), the FRMD8 binding region does not overlap with the sites required for phosphorylation-dependent 14-3-3 binding (Grieve et al., 2017, Cavadas et al., 2017). Consistent with this, the interaction of FRMD8 with iRhom2 was not changed upon PMA stimulation (Fig. 2E). Moreover, an iRhom2 mutant, in which 15 conserved phosphorylation sites have been mutated to alanine (iRhom2^pDEAD^, Fig. S2C) (Grieve et al., 2017), did not abolish the interaction to FRMD8 (Fig. 2E), further demonstrating that the binding of FRMD8 to iRhom2 is independent of the phosphorylation state of iRhom2.

Interestingly, the FRMD8 binding site is absent in a mouse iRhom2 mutant called *curly-bare (cub)*, which lacks residues 1-268 (Hosur et al., 2014, Siggs et al., 2014). Sequence alignment shows that the deletion of 268 amino acids in mouse iRhom2 corresponds to the loss of residues 1-298 in the human protein (Fig. 2C, S2B). Consistent with this mapping data, we found that whereas full-length mouse iRhom2 bound human FRMD8, the *cub* mutant form cannot (Fig. 3E). This failure of FRMD8 binding presumably contributes to the complex defects that underlie the *cub* phenotype (Johnson et al., 2003, Hosur et al., 2014, Siggs et al., 2014).

### FRMD8 protects iRhom2 and ADAM17 from lysosomal degradation

These experiments demonstrate that FRMD8 binds to the cytoplasmic N-terminal region of iRhom2, which has previously been shown to be required to stabilise ADAM17 at the cell surface (Grieve et al., 2017). We therefore tested whether FRMD8 is necessary for iRhom2 to stabilise ADAM17. Treatment of HEK293T wild-type and FRMD8 knockout cells with the lysosomal degradation inhibitors bafilomycin and ammonium chloride restored the mature form of ADAM17 (Fig. 4A; S3A). This result explains the reduced level of mature ADAM17 in FRMD8 knockout cells: it implies that the defect caused by loss of FRMD8 is not a failure of ADAM17 maturation, but instead a failure to stabilise the mature form. In line with this interpretation, the proteasomal inhibitor MG132 had no effect on the stability of mature ADAM17 (Fig. 4A). We conclude that FRMD8 binding to iRhom2 acts to promote ADAM17 function by ensuring its stability after its maturation in the *trans-*Golgi network.

**Figure 4.**
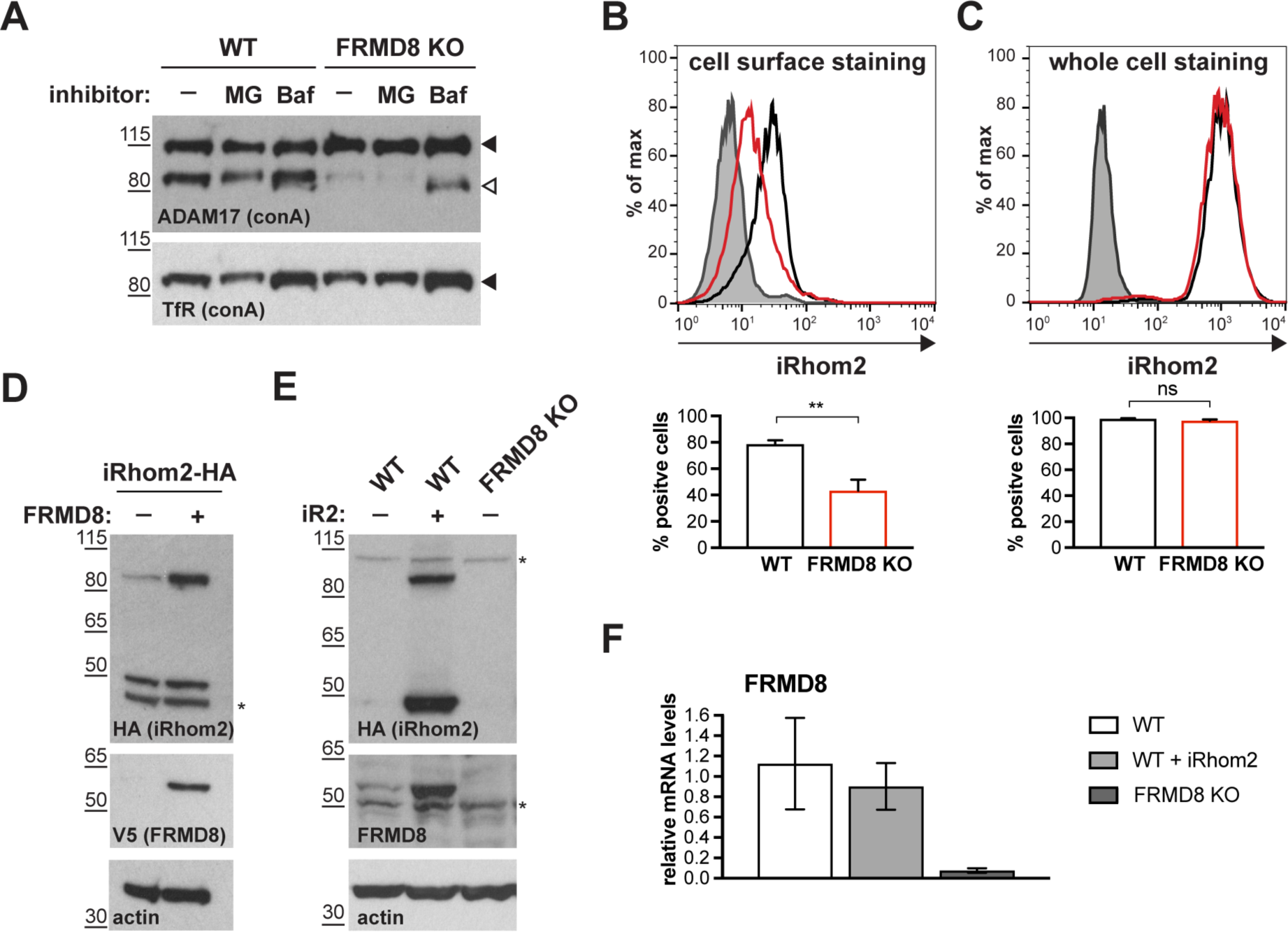
FRMD8 loss leads to the destabilisation of ADAM17 and iRhom2. **A** Cell lysates of wild-type (WT) and FRMD8 knockout (KO) HEK293T cells treated with the solvent DMSO (–), 10 μM MG-132 (MG) or 200 nM bafilomycin A1 (Baf) for 16 h were enriched for glycosylated proteins using concanavalin A (conA) beads and immunoblotted for ADAM17 and transferrin receptor 1 (TfR). TfR was used as a loading control although it is also susceptible to bafilomycin treatment. **B** Unpermeabilised WT (black) and FRMD8 KO HEK293T (red) cells stably expressing human iRhom2-3xHA were immunostained on ice for HA. Wild-type HEK293T cells immunostained for HA served as a negative control (grey). **C** Cells were permeabilised and stained at room temperature with an anti-HA antibody. Immunostaining with the Alexa Fluor 488-coupled secondary antibody served as a control (grey). In (B & C), the flow cytometry graphs shown are one representative experiment out of three experiments. The percentage of positive cells was calculated for each experiment using FlowJo software. Statistical analysis was performed using an unpaired t-test; ns = p-value > 0.05; ** = p-value < 0.01. **D** Lysates of HEK293T cells stably expressing human iRhom2-3xHA and transfected with FRMD8-V5 (where indicated) were analysed by western blot for iRhom2 levels using anti-HA, anti-V5 and anti-actin immunostaining. Nonspecific bands are marked with an asterisk (*). **E** Lysates of WT and FRMD8 KO HEK293T cells stably expressing human iRhom2-3xHA (where indicated) were immunoblotted for HA, FRMD8 and actin. An asterisk (*) marks a nonspecific band. **F** FRMD8 mRNA levels relative to actin mRNA levels were determined by TaqMan PCR in cells used in (E).

These data led us to test the hypothesis that FRMD8 might act as a stabilising factor for the plasma membrane-localised iRhom/ADAM17 sheddase complex that controls the release of growth factors and cytokines. When assaying total levels of exogenously expressed iRhom2, we did not detect any difference between wild-type and FRMD8-deficient cells (Fig. S1C). However, consistent with previous reports (Maney et al., 2015, Grieve et al., 2017), under these experimental conditions most iRhom2 is ER-localised (Fig. S1A) and the cell surface fraction is relatively small. We therefore used cell surface immunostaining of iRhom2 followed by flow cytometry to measure specifically the pool of iRhom2 at the cell surface. In this case the result was clear: in the absence of FRMD8 there was a significant loss of cell surface iRhom2 (Fig. 4B), although the reduction of total iRhom2 levels was not detectable (Fig. 4C). Consistent with our conclusion that although FRMD8 binds continuously to iRhom2, but primarily functions late in the iRhom2/ADAM17 relationship, we detected no defects in the ER iRhom2/immature ADAM17 interaction in FRMD8 knockout cells (Fig. S3B), nor in the trafficking of iRhom2 from the ER to the Golgi (Fig. S3C).

These results show that by binding to iRhom2, FRMD8 stabilises both iRhom2 and mature ADAM17, protecting them from degradation. A more direct demonstration of this stabilising function is provided by overexpressing FRMD8, which leads to increased levels of exogenously expressed iRhom2 (Fig. 4D), as well as iRhom1 (Fig. S3D). Note that the 50 kD N-terminally truncated fragment of iRhoms detected in western blots (Nakagawa et al., 2005, Adrain et al., 2012, Maney et al., 2015) is not stabilised by FRMD8 expression (Fig. 4D, S3D). This iRhom fragment lacks the cytoplasmic tail, and therefore the binding site for FRMD8, so its insensitivity to FRMD8 is consistent with our model. Intriguingly, the stabilisation of iRhom2 and FRMD8 is mutual: overexpression of iRhom2 consistently led to the stabilisation of endogenous FRMD8 protein (Fig. 4E), without affecting FRMD8 mRNA levels (Fig. 4F). This indicates that the iRhom2-FRMD8 interaction leads to mutual stabilisation of both proteins.

To ensure that our conclusion that FRMD8 stabilises iRhoms was not distorted by our use of overexpressed proteins, and in the absence of a usable antibody against human iRhom2, we used CRISPR/Cas9 to insert a triple HA tag into the *RHBDF2* locus to express endogenously C-terminally tagged iRhom2. siRNA-mediated knockdown of iRhom2 confirmed that this editing was successful (Fig. 5A). The cells showed no defect in ADAM17 maturation (Fig. 5A, S3E), indicating that the tagged protein was functional. In these cells FRMD8 overexpression led to an increase in endogenous iRhom2 levels (Fig. 5A); conversely, siRNA knockdown of FRMD8 caused a reduction of iRhom2 protein (Fig. 5B), but no change of iRhom2 mRNA levels (Fig. 5C). Again, the 50 kDa iRhom2 fragment was not affected by FRMD8 levels (Fig. 5A, B). Parenthetically, this is the first reported evidence for the existence of this iRhom fragment endogenously, although its functional significance remains unclear.

**Figure 5.**
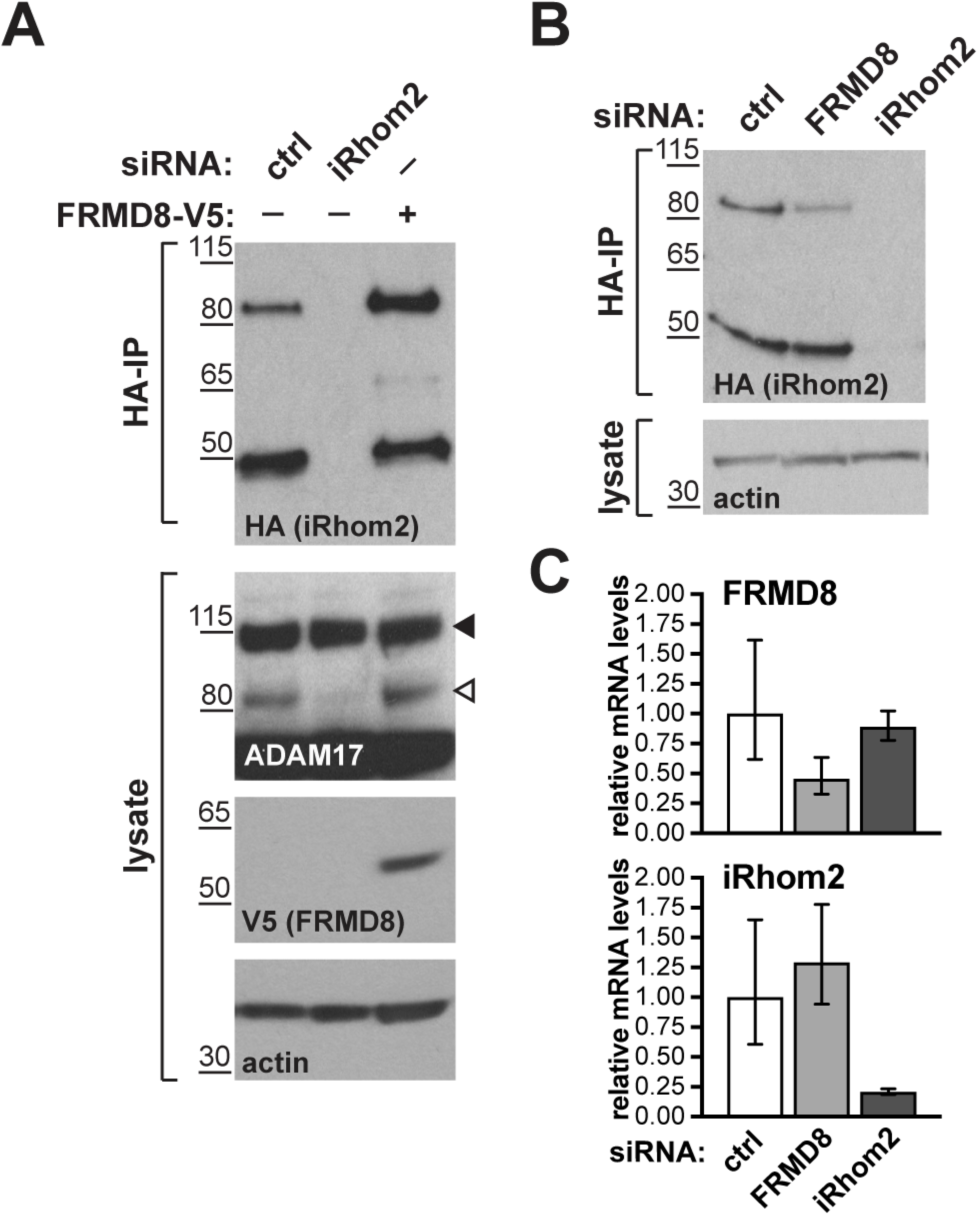
FRMD8 stabilises endogenous iRhom2. **A, B** Levels of endogenously 3xHA tagged iRhom2 were analysed in HEK293T-iRhom2-3xHA cells transfected with FRMD8-V5 plasmid, siRNAs targeting iRhom2, non-targeting siRNA control pool (ctrl) or FRMD8 SMARTpool siRNA. Cell lysates were anti-HA immunoprecipitated (HA-IP) to detect endogenous iRhom2-3xHA levels and immunoblotted using anti-HA antibody. Cell lysates were immunoblotted for ADAM17, V5, and actin. **C** FRMD8 and iRhom2 mRNA levels relative to actin mRNA levels were determined by TaqMan PCR in cells used for the experiment shown in (B) to demonstrate that the destabilisation of endogenous iRhom2 was not induced by a change in iRhom2 mRNA levels.

To summarise our results to this point, we have discovered that by binding to the iRhom2 cytoplasmic N-terminus, FRMD8 is necessary to stabilise the cell surface iRhom2/ADAM17 shedding complex. In the absence of FRMD8, this enzyme complex is degraded by the lysosome. FRMD8 is therefore an essential component of the sheddase complex that releases ADAM17 substrates including cytokines and growth factors.

### FRMD8 binding to iRhom2 is essential for inflammatory signalling in human macrophages

We tested the pathophysiological significance of our conclusions by analysing the consequence of loss of FRMD8 in human macrophages, which release TNFα in response to tissue damage and inflammatory stimuli. To generate mutant human macrophages, we used CRISPR/Cas9 to knock out FRMD8 in an iPSC line that had previously been generated from dermal fibroblasts of a healthy female donor (Fernandes et al., 2016). The FRMD8 knockout and control iPSCs were analysed for deletions in the *FRMD8* gene by PCR (Fig. S4A), and a normal karyotype was confirmed by single nucleotide polymorphism (SNP) analysis (Fig. S4B) before their differentiation into macrophages (Fig. 6A). These mutant macrophages expressed no detectable FRMD8 and, as in the HEK293T cells, showed severely reduced levels of mature ADAM17 (Fig. 6B). When challenged with the inflammatory trigger LPS, TNFα shedding from the cells, as measured by ELISA, was reduced (Fig. 6C). Confirming the expected specificity, the ADAM10 inhibitor GI254023X (GI) had no effect on TNFα release from these cells, whereas GW, an inhibitor of both ADAM10 and ADAM17, further reduced TNFα release (Fig. 6C). Although shedding was inhibited, induction of TNFα expression by LPS was normal in these cells (Fig. S4C). These results demonstrate that our conclusions about the requirement for FRMD8 in ADAM17 function in cell culture models does indeed apply to human macrophages.

**Figure 6.**
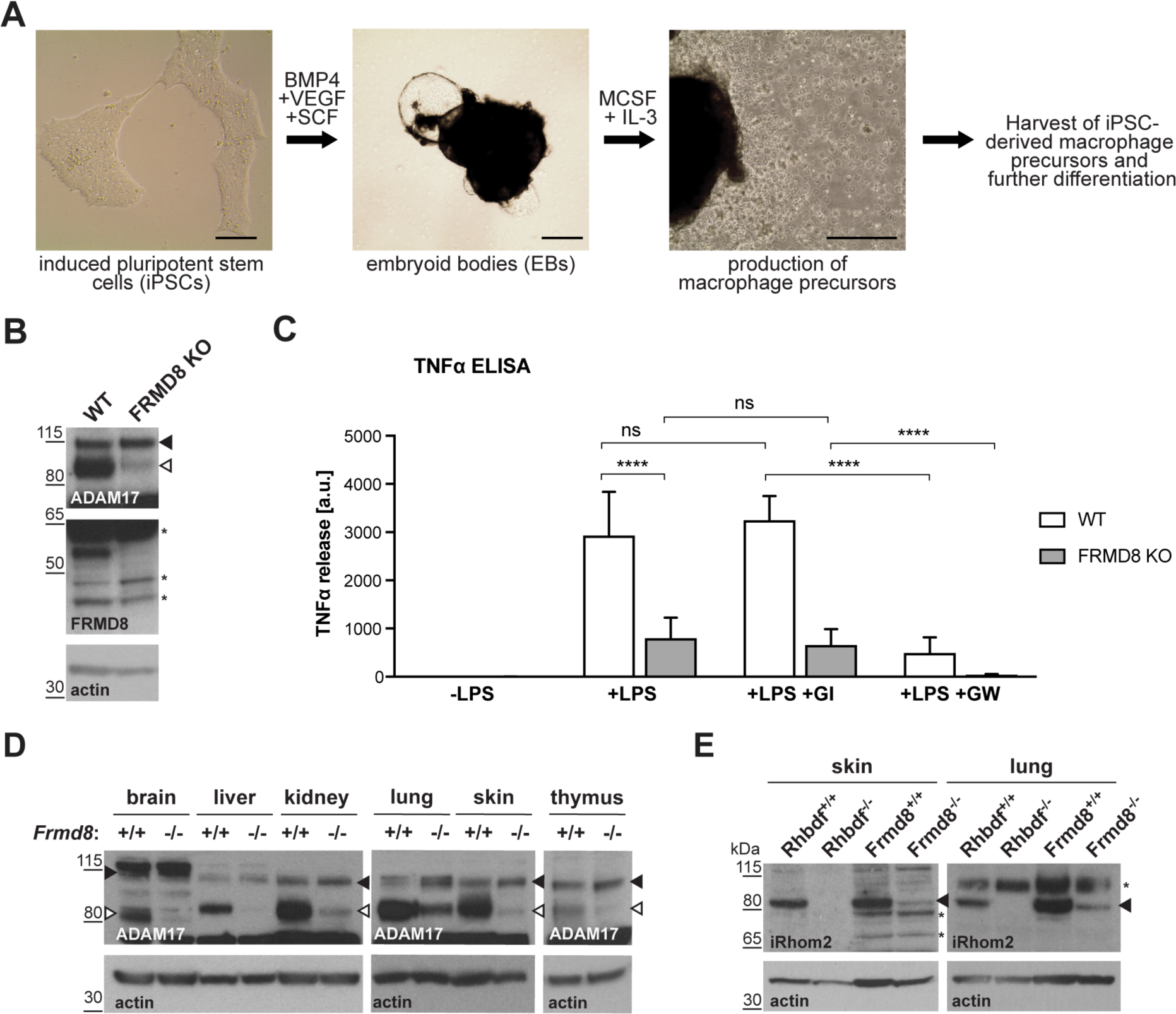
FRMD8 is required for TNFα release in human iPSC-derived macrophages. **A** Schematic representation of the differentiation protocol of iPSCs into macrophages based on (van Wilgenburg et al., 2013). Scale bars: 10 μm. **B** Lysates of iPSC-derived macrophages (on day 7 after harvest from EBs) were immunoblotted for ADAM17, FRMD8, and actin. **C** 25,000 iPSC-derived macrophages were either left unstimulated, stimulated with 50 ng/ml LPS, or with 50 ng/ml LPS and simultaneously with 2 μM GI or 2 μM GW for 4 h. TNFα concentration in the cell supernatants was measured by ELISA and then normalised to the protein concentration in macrophage cell lysates to adjust the cytokine release for potential differences in cell numbers. Each experiment was performed in biological triplicates. Data from three independent experiments were statistically analysed using a Mann-Whitney test; ns = p-value > 0.05; **** = p-value < 0.0001. **D, E** Lysates from tissues derived from *Frmd8^-/-^* or *Rhbdf2^-/-^* and their wild-type littermates were immunoblotted for ADAM17, iRhom2 and actin.

### Loss of FRMD8 in mice highlights its physiological role in stabilising the iRhom/ADAM17 complex

To investigate further the physiological significance of our discovery of the role of FRMD8 in stabilising iRhom/ADAM17 sheddase complexes, we analysed the levels of ADAM17 and iRhom2 in tissues from FRMD8-deficient mice. These mice were generated from embryonic stem (ES) cells from the KOMP Repository, University of California Davis, in which all coding exons (2-11) of the *Frmd8* gene were deleted (Fig. S5A). *Frmd8^-/-^* mice are viable (Fig. S5B) and the knockout was confirmed by western blot (Fig. S5C). Western blot analysis of tissues *Frmd8^-/-^* mice showed that mature ADAM17 levels were reduced in all tissues examined compared to tissues from a wild-type littermates (Fig. 6D). This confirms that FRMD8 controls the level of mature ADAM17 *in vivo*. Of note, there was a major reduction of mature ADAM17 levels in the brain, a tissue in which iRhom2 in almost completely absent but iRhom1 levels are high (Christova et al., 2013, Li et al., 2015). This supports our hypothesis that FRMD8 regulates mature ADAM17 levels through iRhom1 as well as iRhom2. We also tested *in vivo* our conclusion that FRMD8 loss destabilises endogenous iRhoms (Fig. 5B). Using an antibody that we had previously generated against mouse iRhom2 (Adrain et al., 2012), we analysed iRhom2 levels in *Frmd8^+/+^* and *Frmd8^-/-^* mouse tissues. In lung and skin, both tissues with high iRhom2 expression (Christova et al., 2013), we detected a strong decrease of iRhom2 protein levels in *Frmd8^-/-^* compared to wild-type (Fig. 6E). Tissue from *Rhbdf2^-/-^* mice served as a control for the iRhom2 antibody specificity (Fig. 6E). In summary, our experiments in mice confirm the physiological importance of our prior conclusions: FRMD8 is required *in vivo* to regulate the stability of the iRhom/ADAM17 shedding complex, and is therefore a previously unrecognised essential component in regulating cytokine and growth factor signalling.

## Discussion

ADAM17 is the shedding enzyme that is responsible for not only the activation of inflammatory TNFα signalling, but also the release from the cell surface of multiple EGF family growth factors, and other proteins. Its regulation has therefore received much attention, both from the perspective of fundamental cell biology and because of the proven therapeutic significance of blocking TNFα (Monaco et al., 2015). Here we report that FRMD8 is a new component of the regulatory machinery that controls the release of ADAM17 substrates, including TNFα. We identified FRMD8 as a prominent binding partner of iRhoms, which are rhomboid-like proteins that act as regulatory cofactors of ADAM17. Our subsequent experiments demonstrate that although FRMD8 binds to iRhoms throughout their life cycle, its function appears to be confined to the later stages of their role in regulating ADAM17. FRMD8 stabilises the iRhom2/ADAM17 complex at the cell surface, ensuring it is available to shed TNFα and growth factors. We took advantage of iPSC technology to generate human FRMD8 knockout macrophages, allowing us to confirm that the mechanistic conclusions derived mostly from HEK293T cell models were indeed relevant to the human cells that provide the primary inflammatory response. Finally, tissues from FRMD8 knockout mice demonstrate the physiological importance of FRMD8 in a whole organism, and confirm that it stabilises the iRhom/mature ADAM17 complex *in vivo*.

Bringing together all our results, we propose the following model of FRMD8 function in ADAM17-dependent signalling: FRMD8 binds to the cytoplasmic domain of iRhoms throughout the secretory pathway, forming a tripartite complex when iRhoms are also bound to ADAM17. Despite this long-term relationship, we have found no evidence for a functional role for FRMD8 in ER-to-Golgi trafficking or ADAM17 maturation. Instead, FRMD8 acts later, to prevent the endolysosomal degradation of the iRhom/ADAM17 complex (Fig. 7). As we have previously reported, it is this complex that is responsible for shedding ADAM17 substrates including, notably, TNFα. Without FRMD8, iRhoms and mature ADAM17 are destabilised and the cell cannot shed TNFα in response to an inflammatory challenge. Combined with our previous studies (Grieve et al. 2017), this work has changed our perspective on ADAM17, the central enzyme in cytokine and growth factor shedding. Our evidence implies that it would be more appropriate to consider it as the active subunit of a regulatory complex at the cell surface, where iRhoms provide regulatory functions (Maney et al., 2015, Cavadas et al., 2017, Grieve et al., 2017), and FRMD8 maintains the stability of the iRhom/ADAM17 complex post-ADAM17 maturation. It is essential that a pool of the sheddase is available on the cell surface to execute, for example, rapid cytokine release in response to inflammatory signals induced by bacterial infection.

**Figure 7.**
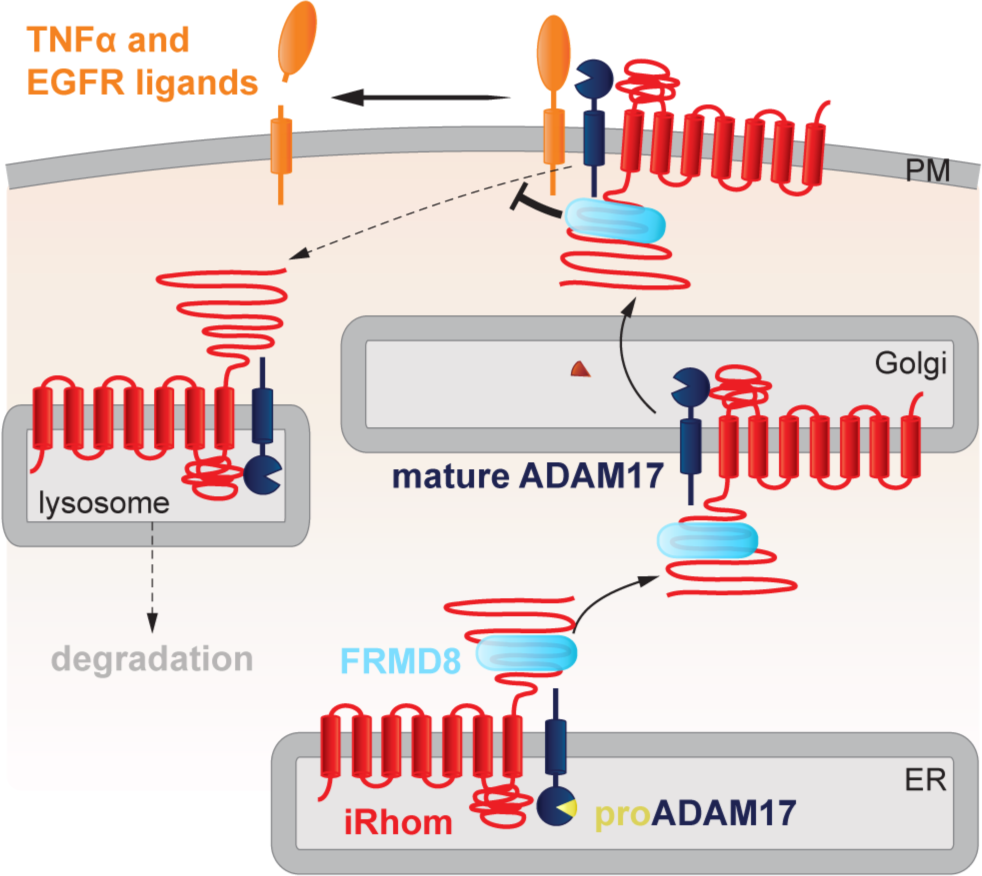
FRMD8 stabilises the iRhom2/ADAM17 sheddase complex. Schematic representation of the role of FRMD8 in the iRhom2/ADAM17 pathway: under wild-type conditions ADAM17 and iRhom2 are stabilised by FRMD8 and thereby protected from degradation through the endolysosmal pathway.

In the only other paper about FRMD8 function, it was reported that FRMD8 (named Bili, after the Drosophila mutation) negatively regulates Wnt signalling by binding to the LRP6 co-receptor, thereby preventing the recruitment of the signal transduction protein axin (Kategaya et al., 2009). Although the signalling event being regulated is different, there is the obvious parallel that in both cases FRMD8 binds to the cytoplasmic tail of a transmembrane protein. In the case of Wnt signalling, this prevents the recruitment of axin; in the case of iRhom function, we do not yet know what the next step in the molecular chain of events is, but the cellular consequence is to prevent recruitment of iRhoms into the endolysosomal degradation system.

Our results extend an important theme to emerge from a number of studies, namely the significance of the cytoplasmic N-terminal region in regulating iRhom function. Several reports indicate that N-terminal mutations cause complex phenotypes that combine aspects of gain and loss of iRhom function, which is consistent with a regulatory function for this region. First, the *cub* mutation, an N-terminal deletion in mouse iRhom2, does not abolish protein function but instead modulates it in complex ways that are still poorly understood. *cub* was described as a gain-of-function mutation that leads to constitutively elevated release of amphiregulin, but is also reported to be defective in releasing TNFα (Hosur et al., 2014). Second, specific point mutations in the N-terminus of human iRhom2 are the cause of a rare genetic disorder called tylosis with oesophageal cancer (TOC) (Blaydon et al., 2012, Saarinen et al., 2012). TOC mutations, as well as truncation of parts of the N-terminus have been reported to enhance the activity of ADAM17 (Maney et al., 2015), leading to the conclusion that parts of the N-terminus have inhibitory functions on ADAM17 function. Third, phosphorylation of specific sites in the iRhom2 N-terminus result in 14-3-3 binding and consequent activation of substrate shedding by associated ADAM17 (Grieve et al., 2017, Cavadas et al., 2017), demonstrating that the N-terminus of iRhom2 also positively regulates ADAM17. Consistent with our current results, we reported previously that iRhom2 lacking the entire N-terminus is not sufficient to support ADAM17-mediated shedding in iRhom1/2-deficient cells, although it can promote ER-to-Golgi trafficking of ADAM17 (Grieve et al., 2017). Complementary to the conclusion that iRhom N-termini are regulatory, the core TMD binding function of iRhoms depends on their membrane-embedded region (Freeman, 2014). A picture therefore begins to emerge of iRhoms having a modular structure, with a core, highly conserved TMD recognition domain in the membrane (and perhaps the lumen), regulated by a more variable N-terminal domain that can integrate cytoplasmic signals.

In light of the growing value of therapeutics that block TNFα signalling, and the wider potential of modulating a wide range of ADAM17 substrates, it is tempting to speculate that the cytoplasmic N-termini of iRhoms could provide potential new drug target opportunities. For example, the limited expression of iRhom2 makes it a theoretically attractive anti-inflammatory target (Issuree et al., 2013, Lichtenthaler, 2013). iRhom2 knockout mice are broadly healthy, beyond defects in TNFα and type I interferon signalling that are only apparent upon challenge by bacterial and viral infections (McIlwain et al., 2012, Luo et al., 2016). Our work now implies that the interface between FRMD8 and iRhoms might be a useful target. This is supported, at least in principle, by our observation that even in cells with complete loss of FRMD8, there is still a low level of mature ADAM17 at the cell surface, and consequently residual TNFα shedding. Even very efficient pharmacological blocking of the FRMD8/iRhom interaction would not, therefore, fully abolish inflammatory responses, potentially reducing side effects. Consistent with this idea, mice with a hypomorphic mutation in ADAM17 show that even only 5% of normal ADAM17 expression is sufficient to rescue many aspects of the loss of function phenotype (Chalaris et al., 2010).

In conclusion, our work demonstrates the cellular and physiological significance of FRMD8 binding to iRhoms, and how this stabilises the iRhom/ADAM17 sheddase complex at the cell surface. It also reinforces the picture that has begun to emerge of ADAM17 not acting alone but instead being supported by at least two other regulatory proteins that act as subunits of what is effectively an enzyme complex. This concept would help to explain how the activity of such a powerful and versatile – and therefore potentially dangerous – shedding enzyme is controlled with necessary precision. The next steps in fully revealing the role of FRMD8 will be to analyse the phenotypic consequences of its loss in mice, which should allow us to understand how the roles of FRMD8 roles in ADAM17 activation, Wnt signalling, and any other potential functions, are integrated. Notwithstanding these physiological questions, the work described here already provides a basis for beginning to investigate the potential of targeting the FRMD8/iRhom interface for modulating the release of ADAM17 substrates.

## Materials and methods

### Molecular cloning

Human UNC93B1, human iRhom2^WT^, iRhom2^Δ100^, iRhom2^Δ200^, iRhom2^Δ300^, iRhom2^Δ382^ were amplified from human UNC93B1 (BC025669.1) and iRhom2 cDNA (NM_024599.2; Origene (SC122961)) by PCR and cloned with an C-terminal 3xHA tag into the lentiviral vector pLEX.puro using Gibson assembly (New England Biolabs) following the manufacturer’s instructions. C-terminal V5-tagged FRMD8 (FRMD8-V5) was amplified from human FRMD8 cDNA (NM_031904; Addgene (SC107202)) by PCR and cloned into pcDNA3.1(+) using Gibson assembly. All constructs were verified by Sanger sequencing (Source Bioscience, Oxford, UK).

### Transfection and transduction of cell lines

Human embryonic kidney (HEK) 293T and MCF-7 cells were cultured in DMEM (Sigma) supplemented with 10% fetal calf serum (FCS) and 1x penicillin-streptomycin (PS) (all Gibco) at 37 °C with 5% CO_2_. Cells were transiently transfected with DNA using FuGENE HD (Promega). Per 10 cm^2^ growth area 4 μl FuGENE HD was added to 1 μg DNA diluted in OptiMEM (Gibco). The transfection mix was incubated for 20 min at RT and added to cells. Protein expression was analysed 48 h - 72 h after transfection. For knockdown experiments, siRNA was transfected using Lipofectamin RNAiMax (Invitrogen) following the manufacturer’s instructions. Per 6 well 50 pmol of FRMD8 SMARTpool siRNA (Dharmacon; siGENOME Human FRMD8 (83786) siRNA; M-018955-01-0010), non-targeting siRNA control pools (Dharmacon; siGENOME D-001206-13-50), RHBDF2 siRNA (Thermo Fisher Scientific; HSS128594 and HSS128595) were used. Protein expression was analysed 72 h after transfection.

HEK293T cell lines stably expressing human UNC93B1-3xHA and human iRhom2-3xHA were generated by lentiviral transduction using the pLEX.puro vector as previously described (Adrain et al., 2012). Cells were selected by adding 2.5 μg/ml puromycin (Gibco).

### Mass spectrometry and data analysis

HEK293T cells expressing human UNC93B1-3xHA (control) and human iRhom2-3xHA were used for anti-HA co-immunoprecipitation and analysed by mass sprectrometry as described previously in (Grieve et al., 2017). Peptides were injected into a nano-flow reversed-phase liquid chromatography coupled to Q Exactive Hybrid Quadrupole-Orbitrap mass spectrometer (Thermo Scientific). The raw data files generated were processed using the MaxQuant (version 1.5.0.35) software, integrated with the Andromeda search engine as described previously (Cox and Mann, 2008, Cox et al., 2011). Differential protein abundance analysis was performed with Perseus (version 1.5.5.3). A two-sample t-test was used to assess the statistical significance of protein abundance fold-changes. P-values were adjusted for multiple hypothesis testing with the Benjamini-Hochberg correction (Hochberg and Benjamini, 1990).

### Co-immunoprecipitation

Cells were washed with ice-cold PBS and then lysed on ice in Trition X-100 lysis buffer (1% Triton X-100, 150 mM NaCl, 50 mM Tris-HCl pH 7.5) supplemented with EDTA-free protease inhibitor mix (Roche) and 10 mM 1,10-Phenanthroline (Sigma). Cell debris were pelleted by centrifugation at 20,000 g at 4°C for 10 min. Proteins were immunoprecipitated by incubation with anti-HA magnetic beads (Thermo Scientific) or anti-V5 magnetic beads (MBL International) for 1 h at 4°C. Beads were washed with Trition X-100 wash buffer (1% Triton X-100, 300 mM NaCl, 50 mM Tris-HCl pH 7.5). Proteins were eluted in 2x LDS buffer (life technologies) supplemented with 50 mM DTT for 10 min at 65°C.

### SDS-PAGE and western blotting

Lysates were mixed with 4x LDS buffer (life technologies) supplemented with 50 mM DTT and denatured for 10 min at 65°C prior to loading on 4-12% Bis-Tris gradient gels run in MOPS running buffer (both Invitrogen). Proteins were transferred to a polyvinylidene difluoride (PVDF) membrane (Millipore) in transfer buffer (Invitrogen). The membrane was blocked in 5% milk-TBST (150 mM NaCl, 10 mM Tris-HCl pH 7.5, 0.05% Tween 20, 5% dry milk powder) and then incubated with the primary antibody: mouse monoclonal anti-β-actin-HRP (Sigma, A3854, 1:5000), rabbit polyclonal anti-ADAM17 (abcam; ab39162; 1:2000), rabbit polyclonal anti-FRMD8 (abcam; ab169933; 1:500), rat monoclonal anti-HA-HRP (Roche, 11867423001, 1:2000), goat polyclonal anti-V5 (Santa Cruz, sc-83849, 1:2000), mouse monoclonal anti-transferrin receptor 1, (Thermo Fisher Scientific, 13-6800, 1:2000), and rabbit polyclonal anti-iRhom2 ((Adrain et al., 2012); 1:500). After three washing steps with TBST (150 mM NaCl, 10 mM Tris-HCl pH 7.5, 0.05% Tween 20), membranes were incubated with the secondary antibody for 1 h at RT using either goat polyclonal anti rabbit-HRP (Sigma, A9169, 1:20000), mouse monoclonal anti-goat-HRP (Santa Cruz, sc-2354, 1:5000) or goat polyclonal anti-mouse-HRP (Santa Cruz, sc-2055, 1:5000).

### CRISPR/Cas9 genome editing in HEK293T cells

For CRISPR/Cas9-mediated knockout of FRMD8 the plasmid pSpCas9(BB)-2A-Puro (pX459; Addgene plasmid #48139) co-expressing the wt *Streptococcus pyogenes* Cas9 and the guide RNA (gRNA) was used. For guide RNA (gRNA) design target sequences with a low chance of off targets were selected using online tools (http://crispr.mit.edu; http://www.sanger.ac.uk/htgt/wge). A gRNA targeting exon 7 (ACCCATAAAACGGCAGCTCGTGG), which is present in all FRMD8 isoforms, was cloned into pX459 resulting in pX459-FRMD8-exon7 plasmid. 1 μg plasmid was transfected into a 6-well of HEK293T cells. Cells were selected with puromycin 48 h after transfection to eliminate non-transfected cells. Single colonies were selected to establish clonal cell lines. Loss of FRMD8 expression was analysed by western blot and quantitative PCR.

CRISPR/Cas9-mediated knock-in of a triple HA tag a homology construct consisting of the triple HA tag flanked at both sides homology arm of approximately 800 bp was cloned into pcDNA3.1(+). The *RHBDF2* locus was targeted in exon 19 in close proximity to the stop codon using gRNA (CCCAGCGGTCAGTGCAGCACCT or CAGCGGTCAGTGCAGCACCTGG) cloned into vector epX459(1.1) (a kind gift from Dr Joey Rieosaame, University of Oxford). HEK293T cells were treated with 200 ng/ml nocodazole (Sigma-Aldrich) for 17 h and then transfected with Cas9/sgRNA plasmid and the pcDNA3.1(+) homology plasmid (0.5 μg each per 6-well). After puromycin selection, single clones were selected to generate clonal cell lines which were tested for the insertion of the tag by PCR.

### mRNA isolation and quantitative RT-PCR

Cells were harvested in PBS and pelleted at 3000 g, 5 min, 4°C. RNA was isolated using the RNeasy kit (Qiagen) and reverse transcribed using the SuperScript VILO cDNA synthesis kit (Invitrogen). Resulting cDNA was used for quantitative PCR (qPCR) using the TaqMan Gene Expression Master Mix (Applied Biosystems) and the following TaqMan probes (all Thermo Fisher Scientific): human ACTB (Hs99999903_m1), human FRMD8 (Hs00607699_mH), human RHBDF2 (Hs00226277_m1), and human TNFα (Hs00174128_m1). qPCR was performed on a StepOnePlus system (Applied Biosystems). Gene expression was normalized to ACTB expression and expressed as relative quantities compared to the corresponding wild-type cell line. Error bars indicate the standard derivation of technical replicates.

### Shedding assay

8 x 10^4^ HEK293T cells were seeded in triplicates per condition into poly-(L)-lysine (PLL)-coated 24-well plates and transfected the next day with 30 ng plasmid DNA encoding Alkaline Phosphatase (AP)-conjugated AREG, HB-EGF or TGFα (received from Prof Carl Blobel). 48 h after transfection, cells were washed with OptiMEM and then incubated with 200 μl phenolred-free OptiMEM (Gibco) containing either 200 nM PMA, the corresponding volume of the solvent (DMSO), or 200 nM PMA and 1 μM GW for 30 min at 37°C. Cell supernatants were collected, the cells were washed in PBS and lysed in 200 μl Trition X-100 lysis buffer. The activity of AP in cell lysates and supernatants was determined by incubating 100 μl AP substrate p-nitrophenyl phosphate (PNPP) (Thermo Scientific) with 100 μl cell lysate or cell supernatant at RT followed by the measurement of the absorption at 405 nm. The percentage of AP-conjugated material released from each well was calculated by dividing the signal from the supernatant by the sum of the signal from lysate and supernatant. The data was expressed as mean of at least three independent experiments, each of which contained three biological replicates per condition.

### Deglycosylation assay

Cells were lysed in Triton X-100 lysis buffer as described above. Lysates were first denatured with Glycoprotein Denaturing Buffer (New England Biolabs) at 65°C for 15 min and then treated with endoglycosidase H (Endo H) or peptide:*N*-glycosidase F (PNGase F) following the manufacturer’s instructions (New England Biolabs).

### Flow cytometry

For ADAM10 and ADAM17 cell surface staining, HEK293T cells were stimulated with 200 nM PMA for 5 min before harvest in PBS. 0.5 x 10^6^ HEK293T cells were washed with ice-cold FACS buffer (0.25% BSA, 0.1 % sodium azide in PBS) and stained with rabbit polyclonal anti-HA antibody (Santa Cruz (sc-805); 0.5 μg diluted in FACS buffer), mouse monoclonal anti-ADAM10 (Biolegend (352702); 4 μg diluted in FACS buffer) or mouse monoclonal anti-ADAM17 (A300E antibody (Yamamoto et al., 2012); 8 μg diluted in FACS buffer) on ice for 45 min. After two washes with FACS buffer, the cells were incubated with Alexa Fluor 488-coupled secondary antibody (Invitrogen (A21202 or A21206); 1:1000 dilution in FACS buffer) on ice for 30 min. Cells were washed twice with ice-cold FACS buffer and then analysed with a BD FACSCalibur (BD Biosciences) and FlowJo software. Cells stained only with the secondary antibody or anti-HA negative cells served as control.

### Culture of human iPSCs

To generate iPSC-derived FRMD8 knockout macrophages, the human iPSC line AH017-13 was used. The AH017-13 line was derived from dermal fibroblasts of healthy donor in the James Martin Stem Cell Facility, University of Oxford as published previously (Fernandes et al., 2016). Donors had given signed informed consent for the derivation of human iPSC lines from skin biopsies and SNP analysis (Ethics Committee: National Health Service, Health Research Authority, NRES Committee South Central, Berkshire, UK (REC 10/H0505/71)). AH017-13 iPSCs were cultured feeder cell-free in mTeSR1 (STEMCELL Technologies) on hESC-qualified geltrex (Gibco). iPSCs were fed daily and routinely passaged with 0.5 mM EDTA, or when required using TrypLE (Gibco) and plated in media containing 10 μmol/l Rho-kinase inhibitor Y-27632 (Abcam).

### Genome editing of iPSCs lines

AH017-13 iPSCs were transfected by electroporation using the Neon Transfection System (Invitrogen). 3 x 10^6^ AH017-13 iPSCs were electroporated (1400 mV, 20 ms, 1 pulse) in a 100 μl tip with 15 μg pX459-FRMD8-exon7 plasmid DNA (endotoxin-free quality), then plated at a density of 4 × 10^5^ cells/cm^2^ and selected 48 h after transfection with 0.25 μg/ml puromycin. After 48 h of selection, surviving cells were plated on a feeder-layer of 4 x 10^6^ irradiated mouse embryonic fibroblasts (MEFs) in 0.1% gelatin-coated 10 cm culture dishes and cultured in hES medium (KnockOut DMEM, 20% KnockOut serum replacement, 2 mM L-Glutamine, 100 μM nonessential amino acids, 50 μM 2-Mercaptoethanol (all Gibco) and 10 ng/mL basic fibroblastic growth factor (bFGF, R&D)). Colonies were manually selected and grown on geltrex in mTeSR1. Clones were analysed by western blot using the anti-FRMD8 antibody, and PCR followed by Sanger sequencing. For PCR DNA was isolated from iPSCs by incubation in DNA isolation buffer (10 mM Tris-HCl (pH 8), 1 mM EDTA, 25 mM NaCl, 200 ug/ml proteinase K added freshly) at 65°C for 30 min. Proteinase K was inactivated at 95°C for 2 min. PCR using Q5 polymerase was performed according to the manufacturer’s instructions (New England Biolabs) using primers FRMD8_fw (TGCAGATCCATGACGAGGA) and FRMD8_rev (GTGCTCGTGACAAGACAC). The PCR product was purified and sequenced using the primer FRMD8_exon7_fw (GCCAGAGTCTCTTTGCTG) for Sanger sequencing (Source Bioscience, Oxford).

### Differentiation of iPSCs into macrophages

AH017-13 wild-type and FRMD8 knockout clones were analysed by Illumina HumanOmniExpress24 single nucleotide polymorphism (SNP) array at the Wellcome Trust Centre for Human Genetics at the University of Oxford and assessed using KaryoStudio software to confirm normal karyotypes before differentiation into macrophages. For this study iPSCs were differentiated into embryoid bodies (EBs) by mechanical lifting of iPSC colonies and differentiated into macrophages as described in (van Wilgenburg et al., 2013). Briefly, iPSCs were grown on a feeder layer of MEFs in hES medium. A dense 10 cm^2^ well of iPSCs was scored into 10x 10 sections using a plastic pipette tip. The resulting 100 patches were lifted with a cell scraper and cell clumps were transferred into a 6-well ultra-low adherence plate (Corning) containing EB formation medium (hES medium supplemented with 50ng/ml BMP4 (Invitrogen), 50ng/ml VEGF (Peprotech) and 20 ng/ml SCF (Miltenyi)) to form EBs. A 50% medium change was performed every second day. On day 5 EBs were harvested.

Approximately 60-80 EBs were transferred into a T75 flask containing factory medium (X-VIVO 15 (Lonza) supplemented with 2 mM L-Glutamine, 50 μM 2-Mercaptoethanol, 100 ng/ml M-CSF and 25 ng/mL IL-3, 100 U/ml penicillin and 100 μg/ml streptomycin (all Gibco)). The EBs were fed weekly with fresh factory medium. After approximately two weeks EBs started to produce non-adherent macrophage precursors, which were harvested from the supernatant of EB cultures through a 70 μM cell strainer. Cells were differentiated into mature adherent macrophages for 7 days in macrophage medium (X-VIVO 15 supplemented with 2 mM L-Glutamine, 100 ng/ml M-CSF, 100 U/ml penicillin and 100 μg/ml streptomycin).

### ELISA

iPSC-derived macrophages were harvested from EB cultures, counted and seeded at 25,000 cells per well into 96-well tissue culture plates in triplicates per condition. Macrophages were cultured in macrophage differentiation medium for 7 days, and then activated with 50 ng/ml LPS (Sigma-Aldrich) in fresh macrophage differentiation medium for 4 h. For inhibitor treatments cells were incubated with 50 ng/ml LPS and 3 μM GW or GI for 4 h. Cell culture supernatants were collected and cleared from cells by centrifugation. TNFα in supernatants was measured by ELISA (Human TNF alpha ELISA Ready-SET-Go, eBioscience (88-7346-86)) according to the manufacturer’s instructions. Macrophages were lysed in Trition X-100 lysis buffer and protein concentration was determined using a BCA assay (Thermo Scientific). The amount of TNFα in the supernatant was normalised to the protein concentration of the corresponding cell lysate to adjust for differences in TNFα release due to cell numbers.

### Mouse work

Commercially available *Frmd8^-/-^* mouse ES cells from KOMP Repository at UC Davis were used to generate *Frmd8^-/-^* mice. The mouse ES cells (C57BL/6NTac strain) were injected into blastocysts of Balb/c mice. Chimeras were bred to C57BL/6 to generate *Frmd8^+/-^* mice that were used for breeding of the colony and the generation of *Frmd8^-/-^* mice. The mouse work was performed under project licenses 80/2584 and 30/2306. Mouse tissues were collected from sacrificed animals and stored on dry ice or at -80°C. Tissues were lysed in Triton X-100 RIPA buffer (1% Triton X-100, 150 mM NaCl, 50 mM Tris-HCl (pH 7.5), 0.1% SDS, 0.5% sodium deoxycholate) supplemented with EDTA-free protease inhibitor mix and 10 mM 1,10-Phenanthroline using a tissue homogeniser (Omni International). Lysates were cleared from cell debris by centrifugation (20,000 g, 4°C, 10 min). Protein concentrations of tissue lysates were determined using a BCA assay.

### Statistical analysis and data presentation

Values are expressed as means of at least three independent experiments with error bars representing the standard deviation. Unpaired, two‐tailed t‐tests were used for statistical analysis. Shedding assays and ELISA data was analysed using a Mann-Whitney test. Flow cytometry blots shown represent one from at least three experiments with similar outcome.

## Acknowledgements

We gratefully acknowledge the support of Oxford’s Advanced Proteomics Facility for our mass spectrometry based proteomic screen and Monika Stegmann for statistical analysis of the results. We also thank Genome Engineering Oxford, specifically Joey Riepsaame and Andrew Bassett, who helped us to design and clone guide RNAs for CRISPR/Cas9 gene editing. We are thankful for the assistance in animal work from the staff of the mouse facility and for support from Elizabeth Robertson, Jonathan Godwin, Angela Moncada Pazos, and Clémence Levet. Immunofluorescent microscopy was performed in Oxford’s Micron imaging facility. We thank members of the Freeman lab for their extensive support throughout this project and their advice on the manuscript. We thank Stefan Düsterhöft and Boris Sieber for providing reagents. This research was supported by the Wellcome Trust to MF (grant number 101035/Z/13/Z). The James Martin Stem Cell Facility has received support from the Wellcome Trust ISSF (121302) and MRC (MC_EX_MR/N50192X/1). UK is supported by the Medical Research Council (award number 1374214) and a Boehringer Ingelheim Fonds PhD fellowship. AG received funding from the European Union’s Horizon 2020 research and innovation programme under the Marie Sklodowska-Curie grant agreement No 659166.

## Author contributions

Conceptualization, U.K., A.G.G., and M.F.; Methodology, U.K., Y.M., and S.A.C.; Investigation, U.K.; Writing, U.K. and M.F.; Funding Acquisition, M.F.

## Supplementary figures

**Supplementary Fig. 1.**
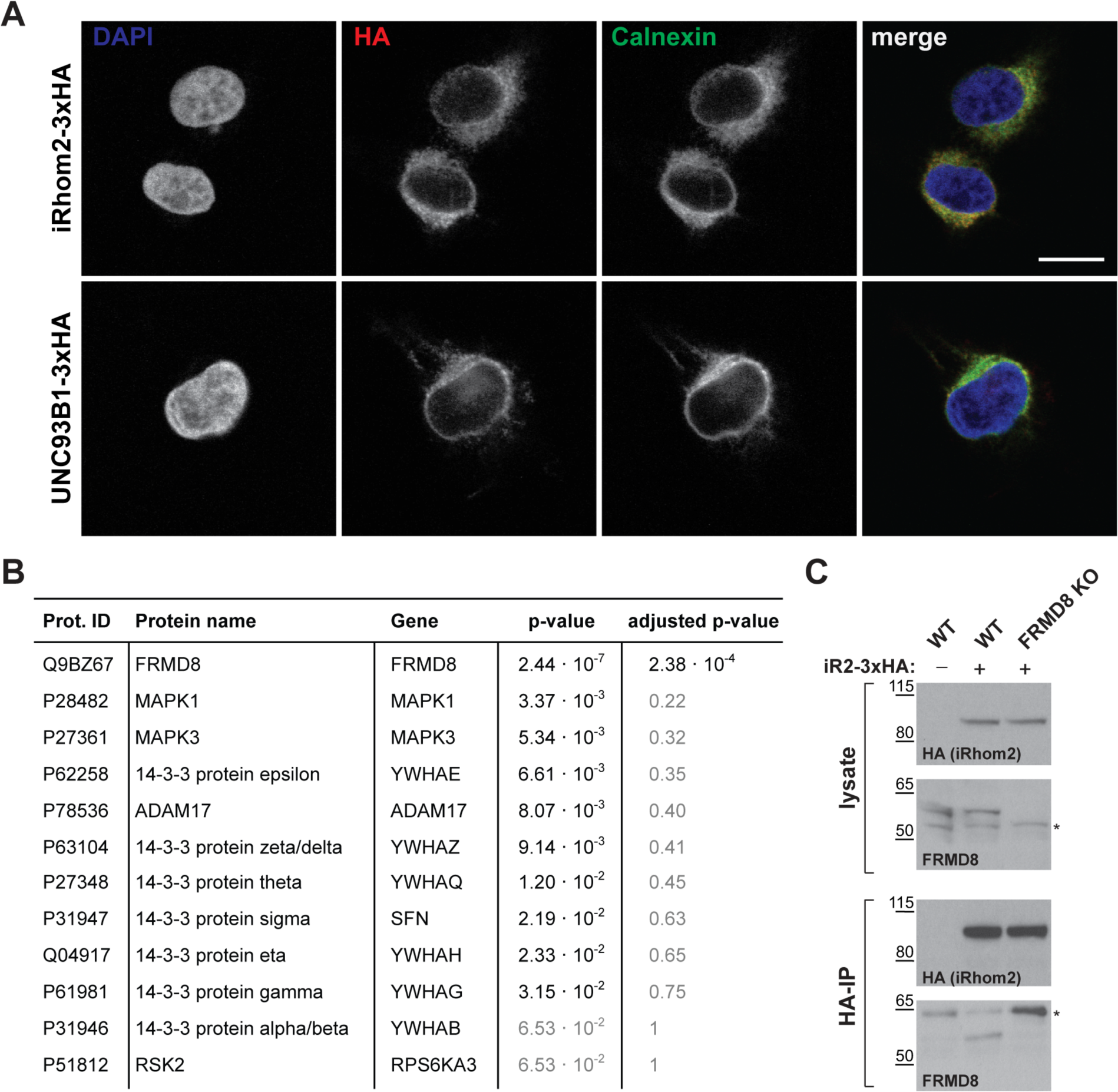
**A** HEK293T cells transiently transfected with human iRhom2-3xHA or UNC93B1-3xHA were stained with 4′,6-diamidino-2-phenylindole (DAPI), blue, to label nuclei, anti-HA to label iRhom2-HA (red), and anti-calnexin to label the ER (green). Scale bar: 10 μm. **B** List of iRhom2 interaction partners identified in the mass spectrometry screen that have either been reported in this study or in previously (Adrain et al., 2012, McIlwain et al., 2012, Grieve et al., 2017). P-values from a two-sample t-test in Perseus are listed with p-values >0.05 written in grey. P-values were adjusted for multiple hypothesis testing with the Benjamini-Hochberg correction and are listed under “adjusted p-values” with p-values > 0.05 written in grey. **C** Lysates and anti-HA immunoprecipitation (HA-IP) from wild-type (WT) and FRMD8 knockout (KO) HEK293T cells stably expressing iRhom2-3xHA (indicated) were immunoblotted for HA and FRMD8. Nonspecific bands are marked with an asterisk.

**Supplementary Fig. 2.**
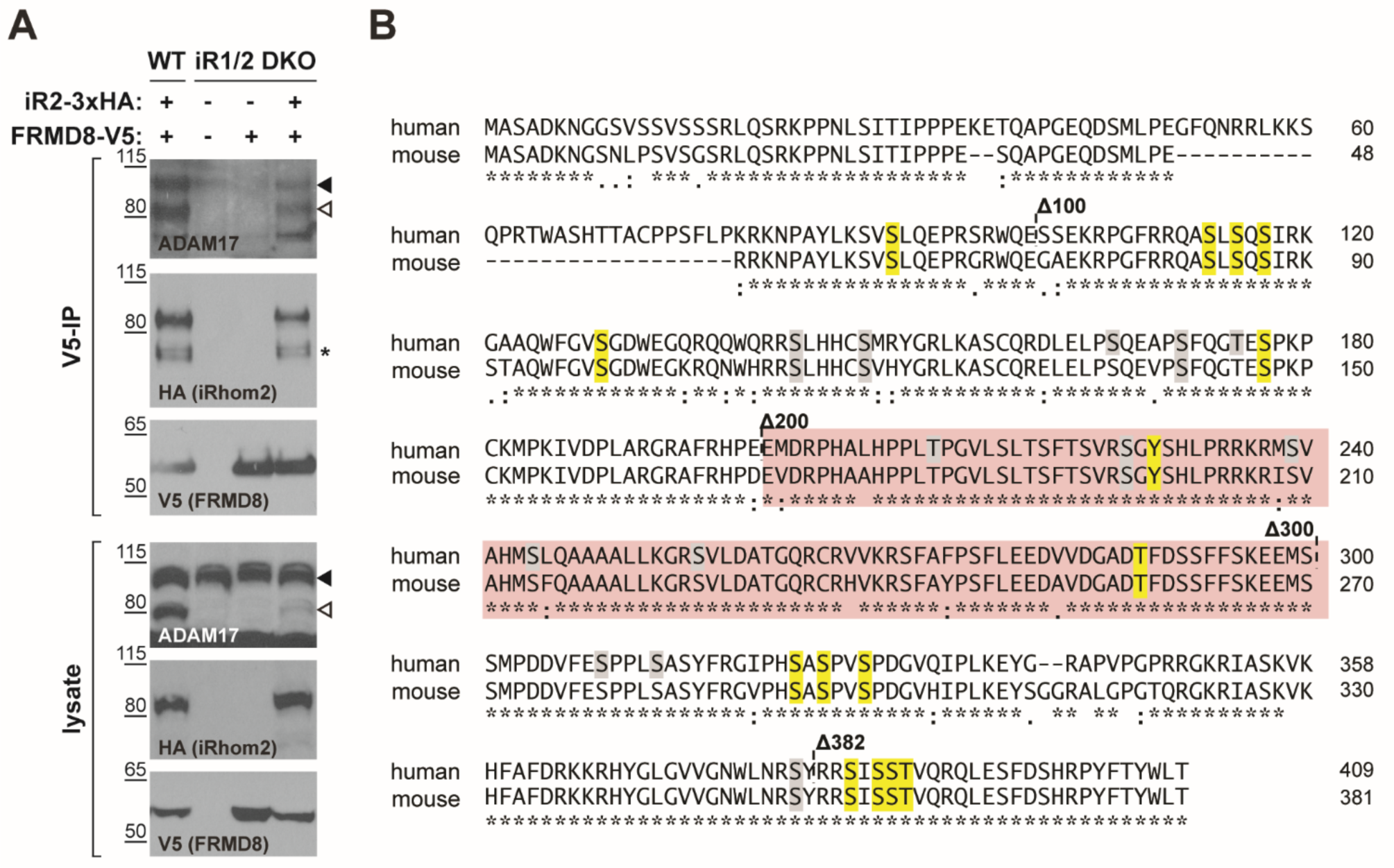
**A** Lysates and anti-V5 immunoprecipitation (V5-IP) from wild-type (WT) and iRhom1/2 double-knockout (DKO) HEK293T cells transiently transfected with iRhom2-3xHA and FRMD8-V5 (where indicated) were immunoblotted for ADAM17, HA and FRMD8. In this and subsequent figures, the pro-and mature form of ADAM17 are indicated with a black and white arrowhead, respectively. An asterisk marks a nonspecific band. **B** Amino acid sequence alignment of human and mouse iRhom2 N-terminal region using Clustal Omega. The region required for FRMD8 binding is highlighted in red. Conserved phosphorylation sites that have been mutated to alanine in theiRhom2^pDEAD^ (Fig. 3E) are marked in yellow. Grey residues indicate additional phosphorylation sites that have been reported on PhosphoSitePlus (www.phosphosite.org). An asterisk (*) indicates positions which have a fully conserved residue, a colon (:) indicates strongly similar properties of the amino acids, and a period (.) indicates weakly similar properties according to the Clustal Omega tool.

**Supplementary Fig. 3.**
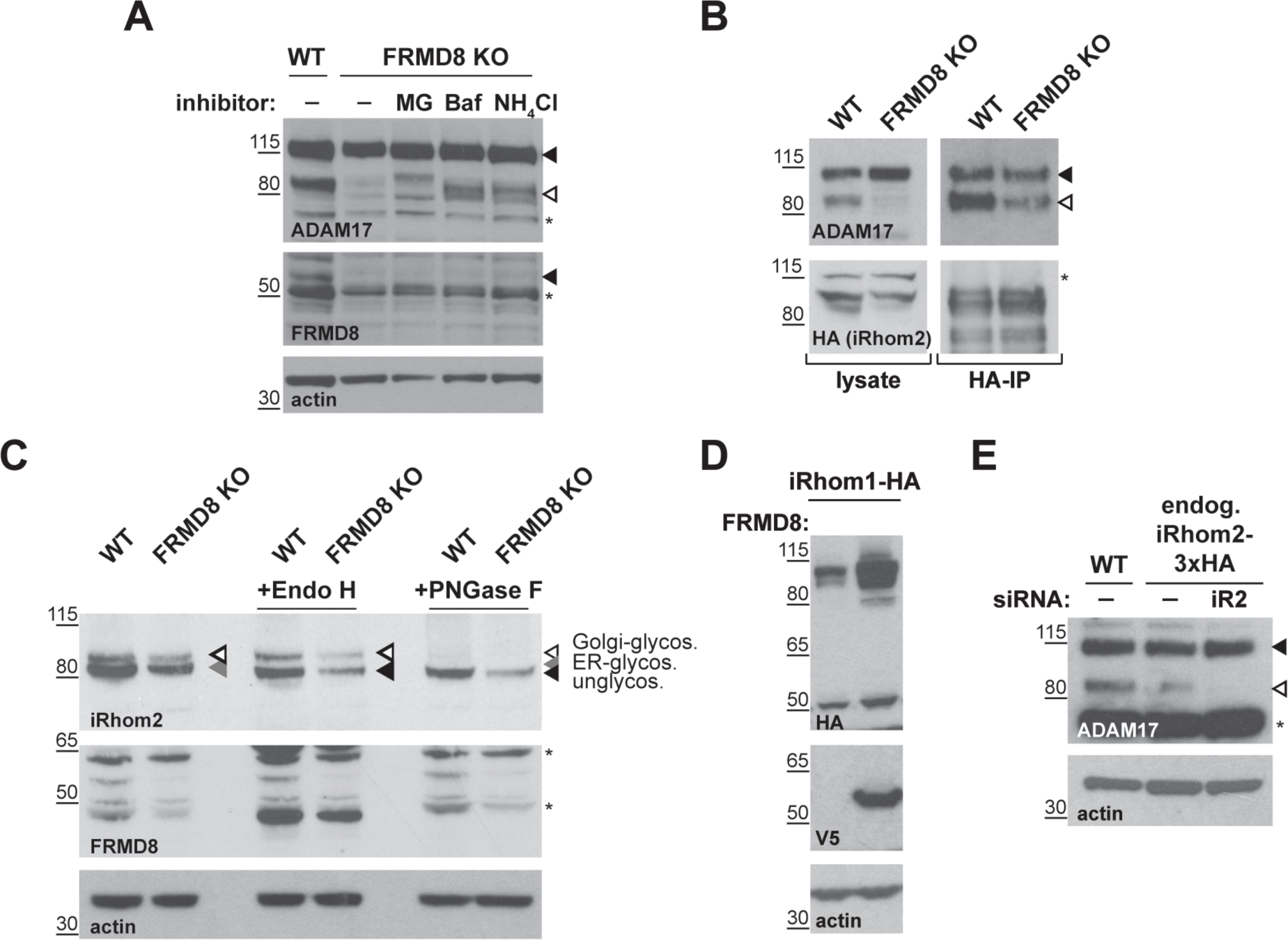
**A** Cell lysates of wild-type (WT) and FRMD8 knockout (KO) HEK293T cells treated with 10 μM MG-132 (MG), 200 nM bafilomycin A1 (Baf) or 50 mM ammonium chloride (NH_4_Cl) for 16 h were immunoblotted for ADAM17, FRMD8, and actin. An asterisk marks a nonspecific band. **B** Lysates of WT and FRMD8 KO HEK293T cells stably expressing human iRhom2-3xHA were anti-HA immunoprecipitated (HA-IP) and immunoblotted for ADAM17 and HA. Nonspecific bands are indicated by an asterisk. **C** N-glycosylation of iRhom2 was analysed using EndoH and PNGase to distinguish ER/*cis-* Golgi (EndoH sensitive) and late Golgi localisation (EndoH resistant). Lysates of WT and FRMD8 KO HEK293T cells transiently transfected with mouse iRhom2-3xHA were deglycosylated with EndoH or PNGase and then immunoblotted for mouse iRhom2, human FRMD8 and actin. An asterisk marks a nonspecific band. **D** Lysates of HEK293T cells stably expressing human iRhom1-3xHA and transfected with FRMD8-V5 (where indicated) were immunoblotted for HA, V5, and actin. **E** Levels of ADAM17 were analysed in HEK293T-iRhom2-3xHA and HEK293T wild-type (WT) cells transfected with siRNAs targeting iRhom2 where indicated. Cell lysates were immunoblotted using an anti-ADAM17 or anti-actin antibody. An asterisk marks a nonspecific band.

**Supplementary Fig. 4.**
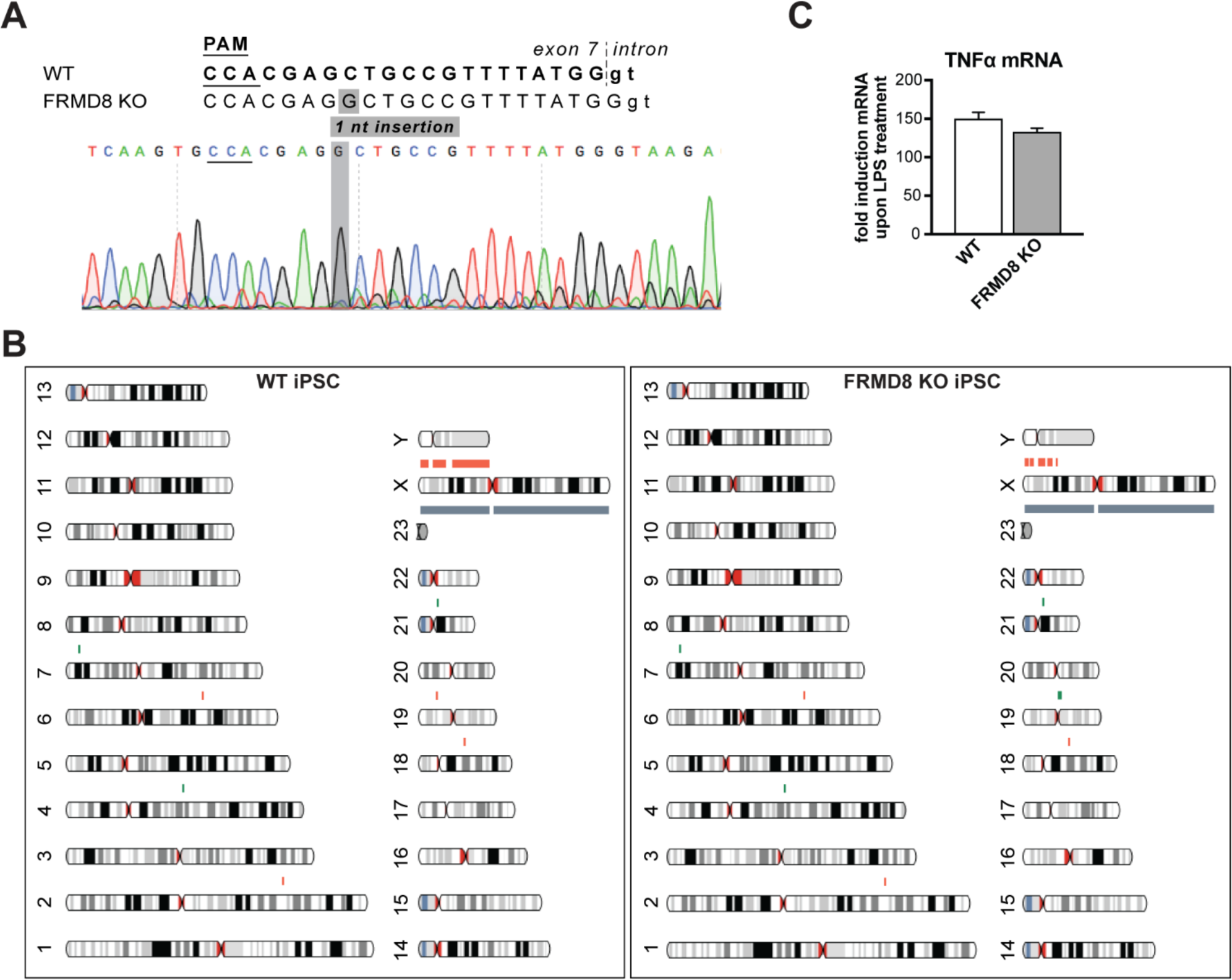
**A** Sequencing of the genomic DNA isolated from clonal FRMD8 KO iPSCs shows a 1-nt insertion. The targeting sequence of the sgRNA is shown in bold (small letters indicate the sequence within the intronic region) with the protospacer adjacent motif (PAM) sequence underlined. **B** Parental wild-type (left) and FRMD8 KO (right) iPSC lines were karyotyped by SNP array. Detected copy number variations are indicated in red (DNA copy number loss in the indicated region) and green (DNA copy number increase). The AH017-13 iPSC line used was derived from a female donor therefore the Y chromosome is marked in red (loss of Y chromosome DNA). **C** TNFα mRNA levels relative to actin mRNA levels were measured by TaqMan PCR in WT and FRMD8 KO iPSC-derived macrophages without stimulation and after stimulation with 100 ng/ml LPS for 4 h. The fold change of TNFα mRNA between unstimulated and stimulated cells is shown.

**Supplementary Fig. 5.**
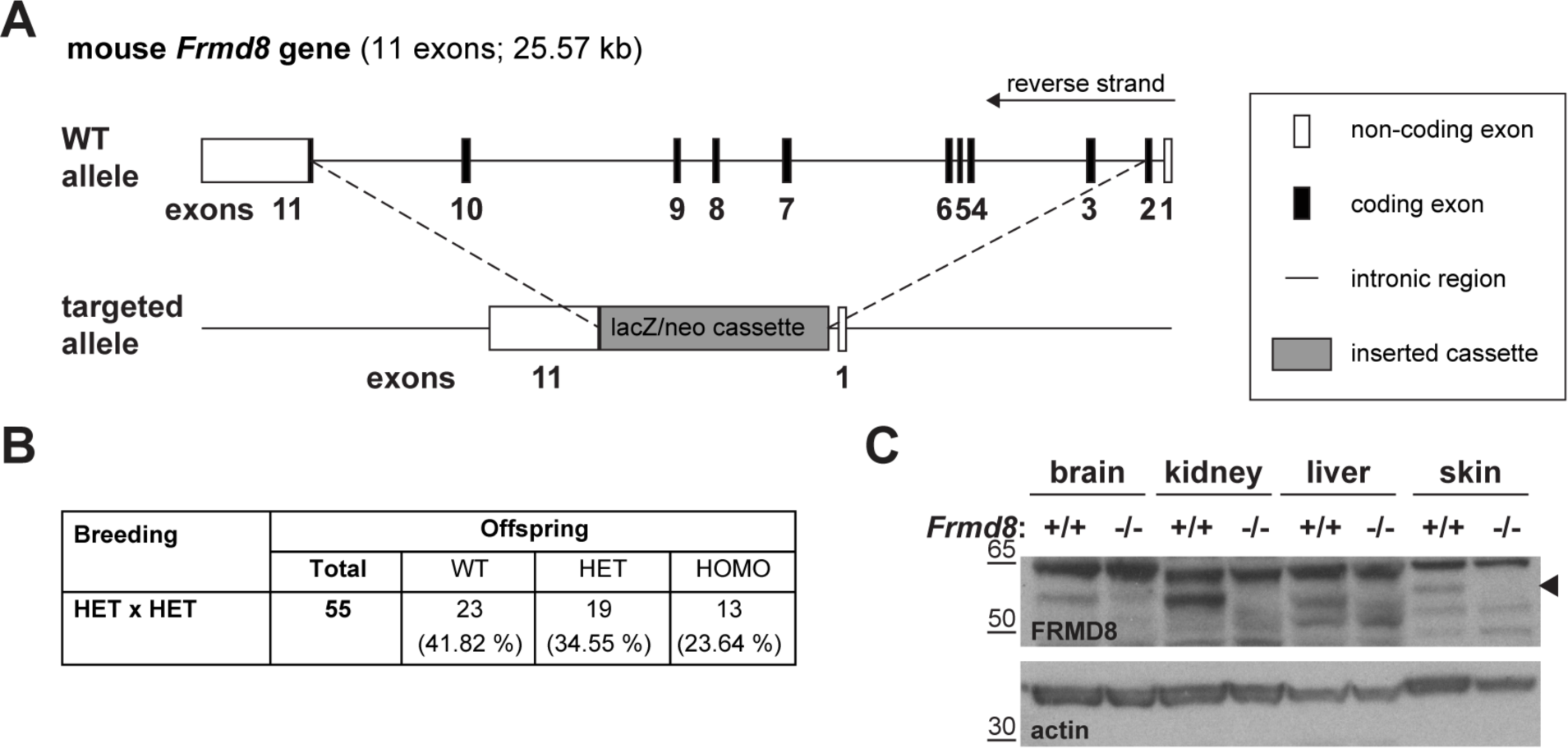
**A** Schematic representation of the insertion of a lacZ/neomycin cassette into the *Frdm8* locus in the ES cells used to generate *Frmd8^-/-^* mice. **B** Off-spring of *Frmd8^+/-^* x *Frmd8^+/-^* (HET x HET) crosses listed by genotype: *Frmd8^+/+^*(WT), *Frmd8^+/-^* (HET), and *Frmd8^-/-^* (KO). **C** Lysates from tissues derived from *Frmd8^-/-^* mice and their wild-type littermate were immunoblotted for FRMD8 and actin to confirm the loss of FRMD8 protein (arrowhead) in the FRMD8-deficient mice.

